# Atomistic simulations of the *E. coli* ribosome provide selection criteria for translationally active substrates

**DOI:** 10.1101/2022.08.13.503842

**Authors:** Zoe L. Watson, Isaac Knudson, Fred R. Ward, Scott J. Miller, Jamie H. D. Cate, Alanna Schepartz, Ara M. Abramyan

## Abstract

As genetic code expansion advances beyond L-α-amino acids to backbone modifications and new polymerization chemistries, the field faces an increasingly broad challenge to discover what the ribosome can accommodate. Although the *E. coli* ribosome tolerates non-L-α-amino acids *in vitro*, few structural insights are available, and the boundary conditions for efficient bond formation are unknown. We describe a 2.1 Å cryo-EM structure of the *E. coli* ribosome containing well-resolved α-amino acid monomers coupled with a computational approach for which energy surface minima produced by metadynamics trend in agreement with established incorporation efficiencies. Reactive monomers across diverse structural classes favor a conformational space characterized by an A-site nucleophile to P-site carbonyl distance of < 4 Å and a Bürgi-Dunitz angle of 90-110°. Monomers whose free energy minima fall outside these regions do not react. Application of this model should accelerate the *in vivo* and *in vitro* ribosomal synthesis and application of sequence-defined, non-peptide heterooligomers.

---

With two exceptions^1,2^, protein chemical space is limited *in vivo* to α-amino and α-hydroxy acids and an otherwise invariant peptide backbone^3–5,1,2^. Not so *in vitro*: under specialized conditions, *E. coli* ribosomes support bond-forming reactions between monomers whose backbones deviate considerably from α-amino acids^6–13^. Yet the yields of peptides containing altered-backbone monomers vary widely, and the products are often detected only by use of autoradiography or mass spectrometry. Here we present a 2.1-Å cryo-EM structure of the *E. coli* 50S ribosomal subunit that visualizes methionine monomers and full-length tRNAs at improved resolution. Using this model, we develop a metadynamics workflow to define the conformational free energy surfaces of structurally and stereochemically diverse altered-backbone monomers within the peptidyl transferase center of the *E. coli* ribosome. Minima in these free energy surfaces clearly differentiate reactive and non-reactive monomers: reactive monomers populate a conformational space characterized by an A-site nucleophile to P-site carbonyl distance of < 4 Å and a Bürgi-Dunitz angle of 90-110°. Monomers whose free energy minima fall outside these regions do not react. This computational workflow is fast, accurate, cost-efficient, and should accelerate the ribosome-promoted biosynthesis of diverse heteroligomers *in vitro* and *in vivo*.

The ribosome is a biological machine that faithfully converts information embodied in one polymer into a different polymer with minimal information loss, synonymous codons notwithstanding. Although the process of mRNA translation enables vast and sophisticated function throughout biology, protein chemical space in extant organisms is limited to about 20 natural α-amino acids and, before post-translational modifications, an invariant peptide backbone. Twenty-plus years of genetic code expansion has broadened protein chemical space to include more than 200 non-natural α-amino acids^5^, but access to alternative backbones *in vivo* is strikingly limited^3,4,1,2^.

Not so *in vitro*, where the concentration of acyl-tRNA can be 50 times higher than possible *in vivo*. Under such conditions, *E. coli* ribosomes support bond-forming reactions between monomers whose backbones deviate considerably from native α-amino acids^6–13^. The structural and electronic diversity of these monomers underscores the importance of proximity in promoting bond-forming reactions within the *E. coli* peptidyl transferase center (PTC) (**Fig. 1a**)^14^. Yet the yields of peptides containing these unusual monomers vary widely, and the products are often detected only by use of autoradiography or mass spectrometry. Given the potential of the ribosome for novel bond-forming chemistry, one must ask: What structural features define reactive monomers? Herein we describe a structurally informed computational workflow designed to identify promising monomers with speed, high accuracy, and low cost.

**Fig. 1.**
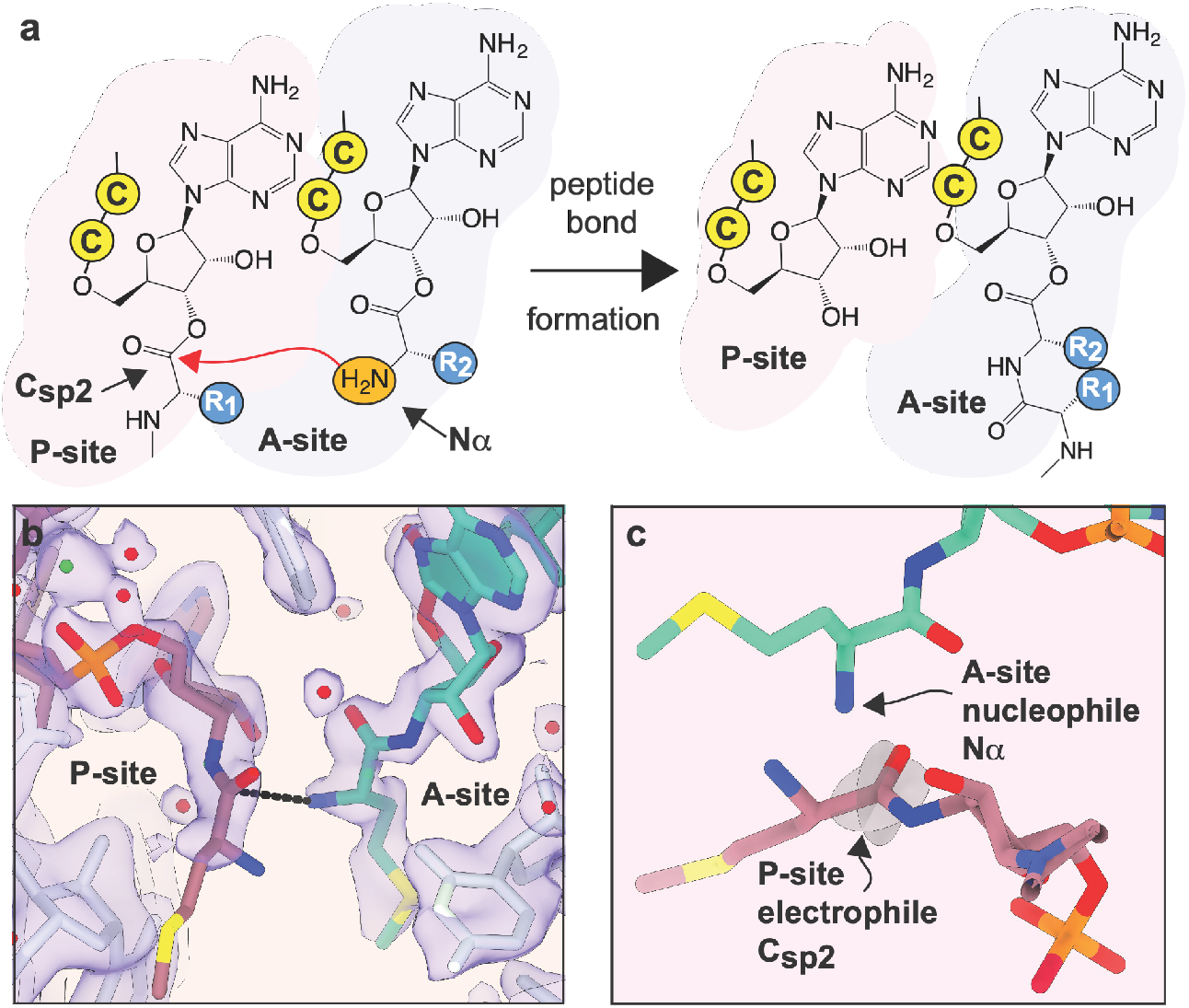
Substrate positioning in the peptidyl transferase center. **a,** Schematic depicting peptide bond formation in the ribosome active site known as the peptidyl transferase center (PTC). The PTC promotes peptide bond formation by positioning the nucleophilic α-amino group (N_α_) of one substrate, the A-site aminoacyl-tRNA, near the electrophilic sp^2^-hybridized carbonyl carbon (C_sp2_) of the second substrate, the P-site peptidyl-tRNA. Attack by N_α_ generates a tetrahedral intermediate that subsequently breaks down to product - a peptidyl-tRNA carrying an additional C-terminal amino acid. Cytidine residues of the tRNAs’ 3’-CCA ends are represented by the letter “C” in a yellow circle. **b,** Cryo-EM density (indigo surface) and our model for Met residues in the PTC. The distance between the A-site (blue) nucleophilic amine and the P-site (purple) carbonyl carbon (3.3 Å) is indicated with a black line. 23S rRNA is shown in white. The metal ion that coordinates the C75-A76 internucleotide phosphate is shown in green. Note that an amide linkage is shown between the A76 and Met residues of each tRNA to reflect the experimental substrates used for structure determination, however, for simulation this was replaced with the ester linkage that would be present during an elongation reaction. The map here was post-processed with a B-factor of −13 Å^2^ and supersampled for smoothness. **c.** Alternative view of the A- and P-site Met residues highlighting the planes of the carbonyl that define the Bürgi-Dunitz (α_BD_) and Flippin-Lodge (α_FL_) angles and the geometry of nucleophilic attack.

We faced three challenges in the design of such a computational schema. First, the size of the ribosome–1.5 MDa in the *E. coli* large ribosomal subunit, not including substrates and solvation–pushes computational methods to their limit^15,16^. As a result, several coarse-grained^17^, normal mode analysis^18^, and elastic network models^19^ combined with simplified ribosome representations have been developed. While these simplifications reduce computational cost and adequately describe large-scale motions, they fail to capture atomistic details. By contrast, density functional theory (DFT) can provide atomistic details, but cannot yet evaluate *de novo* how new monomers engage the PTC in the absence of experimental constraints on orientations in the active site^20^.

The second challenge is that the ribosome is a ribozyme^21^. Most structure-based modeling tools in wide use were developed to study protein enzymes. RNA contains more degrees of freedom than protein, and the polyanionic composition demands accurate, long-range electrostatic and solvation models that include explicit metal ions and water, especially within the PTC^22^. Conformational dynamics alone remains an enormous challenge for computational analyses, with increasing challenges as the complexity of the system increases^23^. While others have used 3D puzzles for RNA structural predictions^24^, physics-based methods such as molecular dynamics (MD) simulations provide a more accurate description of atomistic dynamics of biological systems^25–27^.

The final challenge was the absence of an appropriate high-resolution reference state. The highest resolution structure of the ribosome available at the time of this work is PDB 7K00^28^. This cryo-EM structure includes the 70S *E. coli* ribosome in complex with tRNA and mRNA substrates at 2 Å global resolution. Although local resolution surpassed this value in regions of the large subunit, the PTC itself was less well resolved and the monomers could not be modeled (**Extended Data Fig. 1a**). PDB 6XZ7^29^ is also high-resolution (2.1 Å) but contains product-like tRNAs (**Extended Data Fig. 1b**); PDB 1VY4^22^ contains reactant-like tRNAs but the lower resolution (2.6 Å) obscures details of monomer placement. We concluded that neither PDB 7K00, 6XZ7, nor 1VY4 represent an ideal starting point for simulations to evaluate the conformational landscape of non-L-α-amino acid monomers in the *E. coli* PTC; a structure with well-resolved monomers was required.

Here we present a 2.1-Å cryo-EM structure of the *E. coli* 50S ribosomal subunit that visualizes natural methionine monomers and full-length tRNAs at improved resolution. Using this 50S ribosome model, we developed a metadynamics workflow to define the conformational free energy surfaces of multiple structurally and stereochemically diverse non-α-amino acid monomers within the peptidyl transferase center. Minima in these free energy surfaces clearly differentiate reactive and non-reactive monomers: reactive monomers across all structural classes populate a conformational space characterized by an A-site nucleophile to P-site carbonyl distance of < 4 Å and a Bürgi-Dunitz^30^ angle of 90-110°. Monomers whose free energy minima lie outside a region in which the N_α_–C_sp2_ distance is less than 4 Å, even with an acceptable Bürgi-Dunitz angle, do not react. Metadynamics provided both high accuracy and full conformational sampling of monomers as well the ribosome catalytic center in the explicit solvent and ions environment. This computational workflow is fast, accurate, and relatively cost-efficient. More importantly, it addresses two barriers impeding the ribosome-promoted biosynthesis of diverse heteroligomers. For applications *in vivo*, the MetaD workflow can prioritize monomers for which orthogonal aminoacyl tRNA synthetase variants are needed. For applications *in vitro*, it can identify monomers that are more likely to react within the PTC of wildtype ribosomes and those for which engineered ribosomes are needed. The work reported here should accelerate the *in vivo* ribosomal synthesis of much-sought but not yet achieved sequence-defined, non-peptide heterooligomers and improve the diversity of *in vitro* mRNA display campaigns used in drug discovery.

## Results and Discussion

### Experimental foundational work - improved structure of the *E. coli* PTC

We first established an improved PTC model that retained the high resolution of 7K00^28^ and 6XZ7^29^ but included better-resolved α-amino acid monomers in the A- and P-sites (**Fig. 1b**). We reasoned that the low monomer density seen in 7K00 was due at least in part to aminoacyl-tRNA hydrolysis during grid preparation. To eliminate hydrolysis, we acylated 3’-amino tRNA^fMet^ with Met and *E. coli* MetRS^31^. With this hydrolysis-resistant substrate we obtained a 2.1 Å resolution cryo-EM structure of the *E. coli* 50S subunit from 70S complexes containing well-resolved Met-NH-tRNA^fMet^ in both the A and P sites (**Extended Data Fig. 2**). During data processing, a classification approach more involved than our approach for PDB 7K00^28^ was used to determine a balance between maximal particle inclusion for higher global resolution and more discriminating classification beyond simple tRNA occupancy (see **Methods** and **Extended Data Fig. 3**).

**Fig 2.**
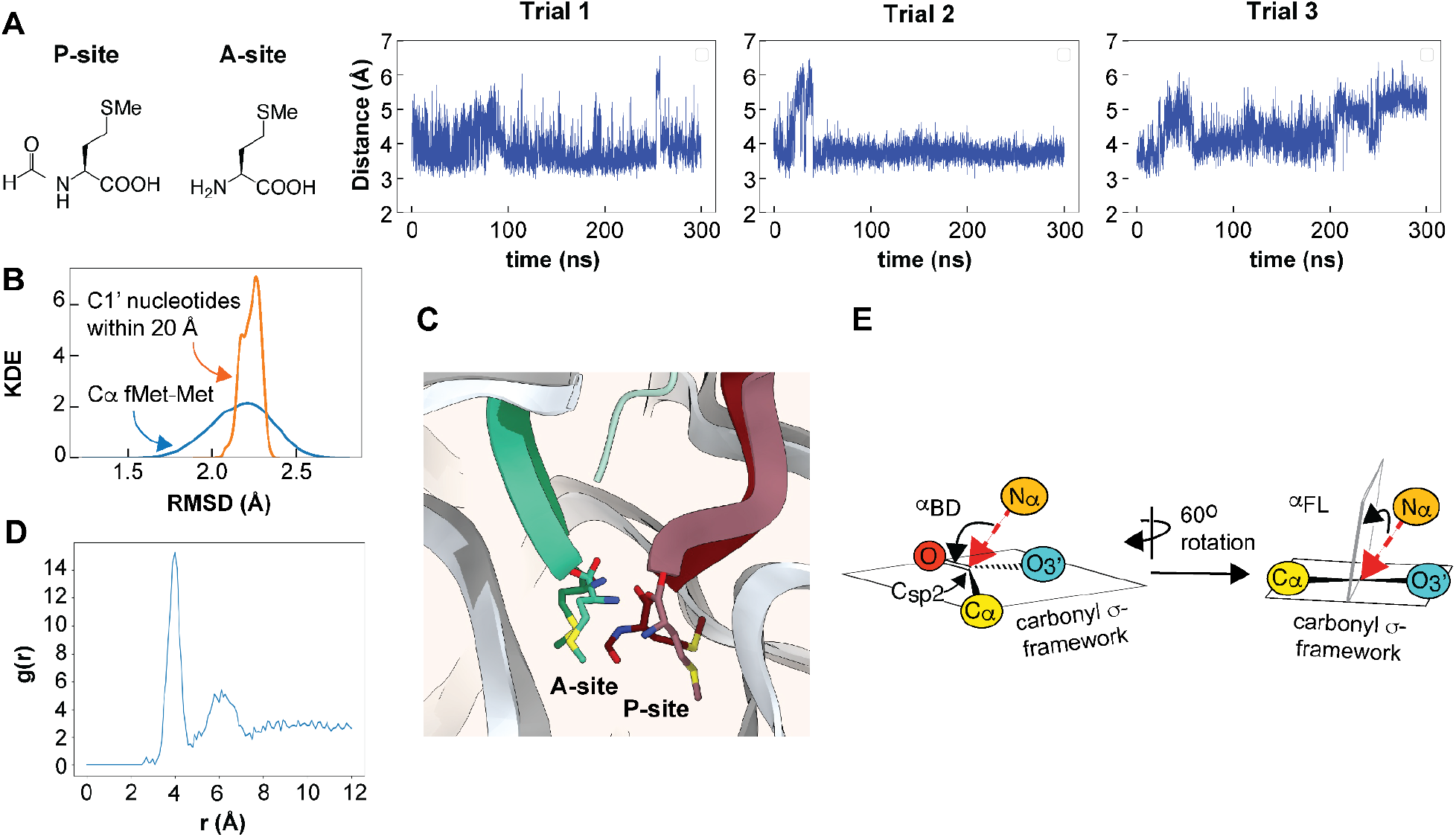
Molecular dynamics simulations of the input fMet-Met ribosome structural model. **a,** Trajectories illustrating evolution of the distance between the A-site Met nucleophile (N_α_) and the P-site fMet carbonyl electrophile (C_sp2_) over 300 ns within the 30 Å Reduced Ribosome Model (RRM). **b,** Kernel Density Estimation (KDE) of the root-mean-square deviation of either the C1’ nucleotides within the internal (non-fixed) 20 Å of the RRM (orange) or the C_α_ carbons of Met and fMet (blue) over the course of the simulation. **c,** Close-up of the bond-forming region of the PTC in the 2.1 Å cryo-EM model reported here (lighter shades) and a representative pose (see Methods) of the simulation (darker shades) illustrating the relative positions of the Met and fMet monomers in the A and P sites, respectively. **d,** Radial distribution function indicates that the first and second solvation shells for the K^+^ and Mg^2+^ ions are within ~4 and 6 Å of the P atoms of residues C75 and A76, respectively. **e,** Illustration of the Bürgi–Dunitz angle (α_BD_), which specifies the angle between the entering nucleophile and the C_sp2_-O double bond and the Flippin-Lodge angle (α_FL_), which specifies the offset of the attack angle from the plane orthogonal to that defined by the carbonyl and the two adjacent substituents.

**Fig. 3.**
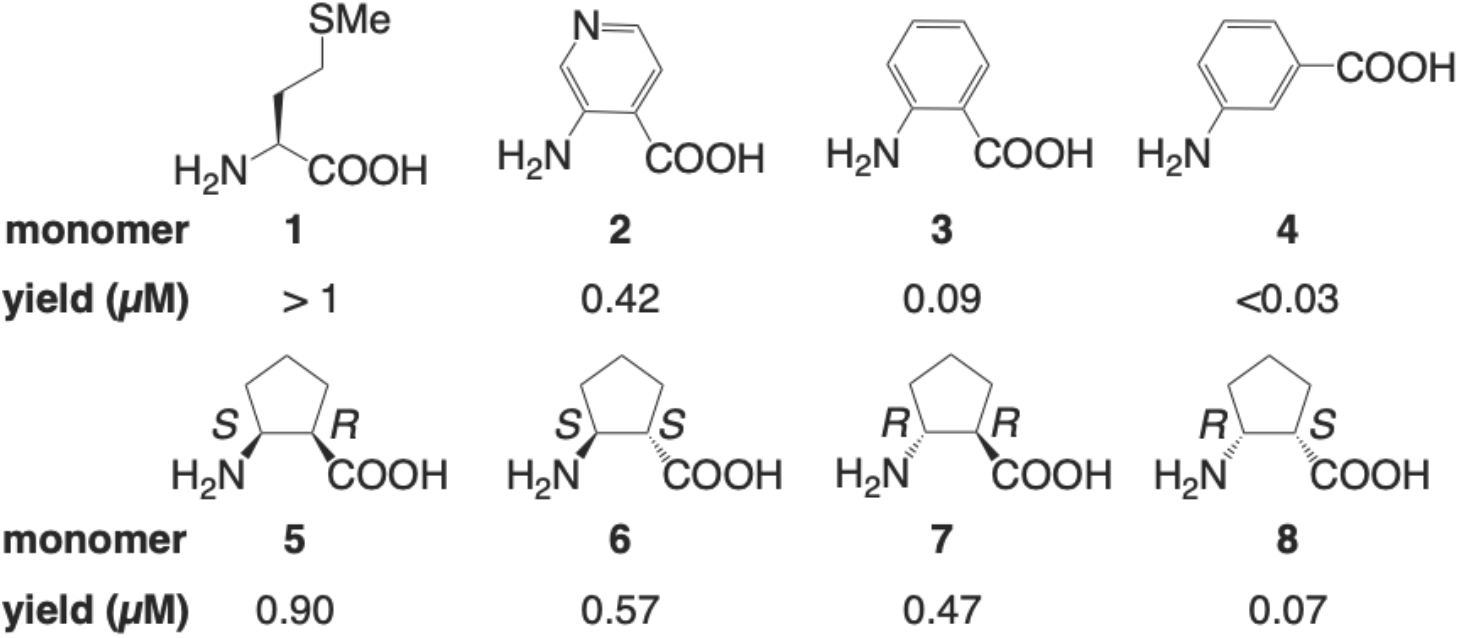
Chemical structures of monomers tested in MD and metadynamics simulations. Structures of aramid and cyclic β^2,3^-2-aminocyclopentane-1-carboxylic acid (ACPC) monomers and data from *in vitro* translation (IVT) assays are shown. The yield shown represents the relative efficiency of translation of DNA template fMet-Trp-Lys-Lys-Trp-Lys-Lys-Trp-Lys-X-Gly-Asp-Tyr-Lys-Asp-Asp-Asp-Asp-Lys (for aramids) or fMet-Trp-Lys-Lys-Trp-Lys-Lys-Trp-Lys-Phe-X-Gly-Asp-Tyr-Lys-Asp-Asp-Asp-Asp-Lys (for ACPC monomers). IVT reaction conditions were comparable within each monomer class, as was the efficiency of tRNA acylation. The value for Met was estimated at 2.5-times the efficiency of Ala.

Local features of the PTC in the current map (**Fig. 1b**) are substantially improved from PDB 7K00 despite lower 2.1 Å global resolution of the large subunit. This resolution was sufficient to improve the modeling of ordered water molecules, ions, and polyamines in the PTC. Density for key bases U2506 and U2585 were modeled suboptimally in 7K00 due to poor density (**Extended Data Fig. 1a**), while in the current structure they have improved density best modeled in the favored *anti* conformation in agreement with PDBs 6XZ7 and 1VY4 (**Extended Data Fig. 4b,c**). The new map also enabled the modeling of residues 2-7 of bL27, the only ribosomal protein that is proximal to the tRNA CCA ends (**Extended Data Fig. 4a**). Importantly for the simulations to follow, residues C75 and A76 in each acyl tRNA are well resolved, with clear positioning of both the ribose and phosphate backbone as well as the amide linkages between Met and each A76 ribose (**Fig. 1b**). Beyond the amide linkage, the entirety of the A-site Met can be modeled. By contrast, the side chain and amine group of the P-site Met are not well resolved. However, the phosphate-ribose backbone of A76 in the P site is more clearly resolved than in previous models, resulting in a relative change in position (**Extended Data Fig. 1c**). In the model, the distance from the nucleophilic A-site amine N_α_ to the P-site carbonyl carbon C_sp2_ is 3.3 Å (**Fig. 1b** and **c**). The ability to visualize these elements is especially important for simulations, in which remaining details of the residues themselves would be substituted *in silico*.

**Fig. 4.**
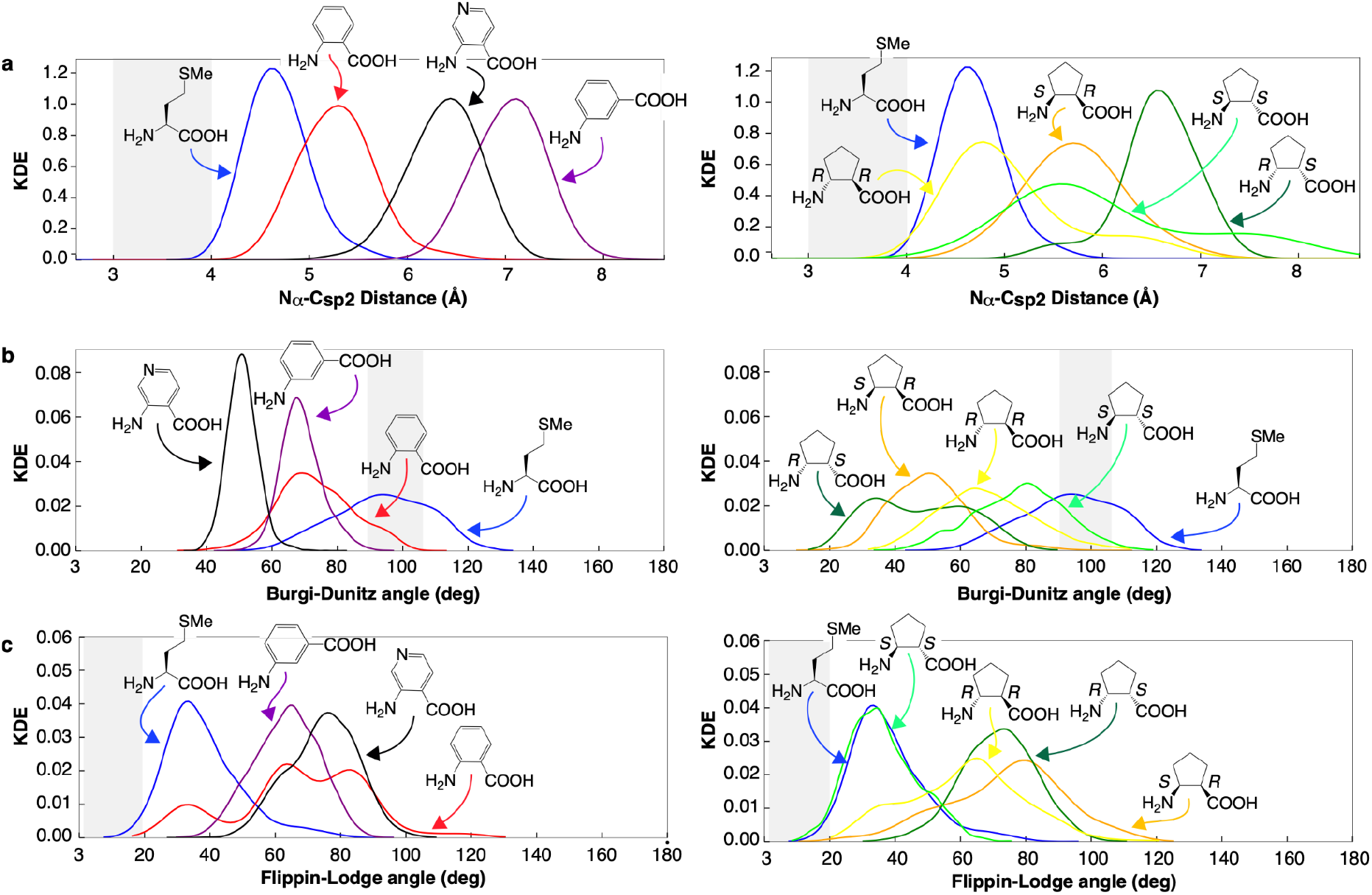
Kernel Density Estimation (KDE) of the three geometric measurements from the MD simulations. **a,** N_α_ – C_sp2_ distance; **b,** Bürgi-Dunitz angle; and **c,** Flippin-Lodge angle for aramids (left) and β^2,3^-amino acid 2-aminocyclopentane-1-carboxylic acid (right) monomers within the RRM over the course of the 300 ns simulation. The gray zones highlight N_α_ – C_sp2_ distances of between 3 and 4 Å, and values of α_BD_ and α_FL_ of between 90° and 105° and 5° to 20°, respectively.

### Computational preliminaries - establishing and validating a test system

It is a well-established approximation, when judiciously applied, that truncation and/or constraining of distant atoms of the ribosome can reduce computational cost while still providing insight about conformational dynamics within the PTC^32–34^. We thus truncated the cryo-EM model reported here to include only those residues within 30 Å of both the A- and P-site Met; this model was solvated in a water box and a physiological concentration of KCl was added. As the simulation environment was set up with preliminary coordinates from the cryo-EM structure which did not yet contain ions or spermidine ligands, ordered Mg^2+^ ions from PDB 7K00 were rigid-body docked into the refined PTC coordinates. This truncation decreased the system size to ~88,000 atoms, including solvent, salt, and counterions. This model is referred to as RRM, for Reduced Ribosome Model.

First we set out to ensure that the RRM was stable during MD simulations and would recapitulate the conformations of the A- and P-site monomers visualized by cryo-EM. We acylated the P-site Met with an *N*-formyl group *in silico* to serve as a more physiological initiator monomer, and positional restraints were placed on the Cα of every ribosomal protein and the C1’ of every nucleotide within the outer 10 Å of the 30 Å model. Test MD simulations (300 ns each) were run in triplicate - each with a different starting velocity to evaluate statistical uncertainties and improve sampling (**Fig. 2a**). As an initial validation step, we examined backbone atom fluctuations during the three 300 ns simulations with respect to the original cryo-EM structure. We evaluated both overall model stability (the RMSD of every unrestrained C1’ atom from its position in the cryo-EM structure) and monomer stability within the PTC (the RMSD of the Cα atoms of Met and fMet) (**Fig. 2b**). Over the course of the simulation, the overall model RMSD was centered at 2.2 Å with a narrow distribution; stability within the PTC also averaged 2.2 Å, but with a slightly wider distribution. These metrics provide confidence that the overall RRM conformation and Met/fMet fluctuations remain stable (**Fig 2c**).

As a further test of the validity of this model system, we examined the distribution of cations in the vicinity of the P-site tRNA 3’-CCA end. In the high-resolution structure determined here - after modeling of ions in the PTC was completed - and in the *T. thermophilus* structure (PDB 1VY4)^22^, a cation coordinates the phosphate group linking C75 and A76 of the P-site tRNA (**Fig. 1b**). Since the tRNA CCA ends were not well-positioned in 7K00^28^, this Mg^2+^ ion was not included in the initial RRM. Notably, in test MD simulations, which included K^+^ and Mg^2+^ ions, strong cation positioning close to the P-site tRNA C75 and A76 phosphates is observed. The radial distribution functions (RDF) for K^+^ and Mg^2+^ ions show ion to phosphorus distances consistent with phosphate coordination during the time course (**Fig. 2d**).

We next examined the geometric relationship between the A- and P-site monomers that react to promote peptide bond formation. This geometric relationship is described by three metrics. The first is the distance between the candidate nucleophilic α-amine of the A-site monomer (N_α_) and the candidate electrophile, the ester carbonyl carbon of the P-site monomer (C_sp2_). This distance is 3.3 Å in the 2.1 Å resolution cryo-EM structure reported here; it averaged 4.0 ± 0.3 Å throughout the three simulations. The second and third metrics are angles that define the approach of the nucleophile relative to the carbonyl (**Fig. 2e**). The Bürgi–Dunitz angle (α_BD_)^30^ specifies the angle between the entering nucleophilic N_α_ and the C_sp2_-O double bond. Values^30,35^ for α_BD_ vary from ~105° to ~90°. The Flippin-Lodge angle (α_FL_)^36,37^ specifies the offset of the attack angle from the plane orthogonal to that defined by the carbonyl and the two adjacent substituents and is smaller, often < 20°. The values of α_BD_ and α_FL_ derived from the cryo-EM structure reported here are 98° and 19°, respectively, also in line with canonical data and predictions. The average values of α_BD_ and α_FL_ derived from the simulations are centered at 94° ± 15° and 38° ± 12°, respectively.

### Combined experimental and computational evaluation - two test sets of non-L-α-amino acid monomers

With a validated workflow in hand, we evaluated the intra-PTC conformational landscapes of two sets of non-L-α-amino acid monomers (**Fig. 3**). These monomers exhibit differences in overall structure, stereochemistry, and basicity of the nucleophilic atom (predicted K_a_ values span 7 orders of magnitude). All acylate a common tRNA in high yield in Flexizyme^38^-promoted reactions, yet the acylated tRNAs that result differ significantly in translational efficiency *in vitro* using *E. coli* ribosomes. The first monomer set comprises a trio of aminobenzoic acid derivatives (**2-4**) that introduce extended, sp^2^-hybridized aromatic backbones into translated polypeptides. Of these three monomers, *ortho*-substituted pyridine **2** is the most reactive, with a translation yield at least 4-fold higher than reactions of tRNA acylated with *ortho*-amino benzoic acid **3**^12^. The translation yield using *meta*-amino benzoic acid **4** was almost undetectable. The second monomer set comprises all four stereoisomers of the cyclic β^2,3^-amino acid 2-aminocyclopentane-1-carboxylic acid (**5**-**8**)^9^ studied extensively as foldamers^39^. Of these four monomers, stereoisomer **5** is the most reactive; the translation yield was more than 10-fold higher than that of enantiomer **8**. The translational efficiencies of the enantiomeric pair **6** and **7** were moderate.

### Incorporation yield is not predicted by average distance between nucleophilic and electrophilic atoms in MD simulations

Next we evaluated in triplicate the evolution of each non-L-α-amino acid monomer over 300 ns within the PTC of the 30 Å RRM. We evaluated changes in the distance between N_α_ and C_sp2_ as well as the values of α_BD_ and α_FL_^36,37^ (**Fig. 4** and **Extended Data Fig. 5**). A plot of the average kernel density estimate (KDE) as a function of all three metrics spans a wide range for both monomer sets (**Fig. 4a**). As described above, simulation of the native fMet-Met pair reproduces the expected values for the N_α_ – C_sp2_ distance (4.0 ± 0.3 Å) and reasonable values of both α_BD_ (94° ± 15°) and α_FL_ (38° ± 12°). However, the midpoint N_α_–C_sp2_ distance for non-L-α-amino acid monomers **2-8** does not correlate with reactivity. For aramids **2-4**, although the midpoint N_α_–C_sp2_ distance for *ortho*-aramid **3** (4.7 Å) is smaller than *meta*-aramid **4** (6.4 Å), it is also smaller than pyridine analog **2** (5.7 Å) which is by far the most reactive. In the case of cyclic β^2,3^-amino acids, reactivity again fails to track with midpoint N_α_–C_sp2_ distance. Reactivity tracks in the order **5** > **6** ~ **7** >> **8**, whereas the the midpoint N_α_–C_sp2_ distance tracks in the order **7** < **5** ~ **6** < **8**. Similar conclusions can be drawn when α_BD_ and α_FL_ are considered in place of the midpoint N_α_–C_sp2_ distance (**Fig. 4b,c** and **Extended Data Fig. 6**).

### Metadynamics provide more thorough sampling of the monomer-dependent conformational landscape within the PTC

Based on the lack of correlation discussed above, we concluded that in the absence of experimentally determined starting structures for non-L-α-amino acid monomers, the *in silico* assembled starting conformations could be trapped in local energy wells and likely did not explore the entirety of conformational space during the course of the MD simulation. This problem is common in MD simulations of biomolecular systems - sometimes the sampling over a given time scale is insufficient, other times the system may be trapped at a local energy minimum and diffuse slowly, other times both of these scenarios occur simultaneously^40,25,41^. To achieve a more thorough sampling of the monomer-dependent conformational landscape within the PTC, we turned to metadynamics^42^ which has been utilized extensively in sampling of biomolecular simulations^43^. Metadynamics improves sampling by introducing an additional force (bias potential) on a chosen number and types of degrees of freedom, known as collective variables (CVs). For practical purposes it is important to limit the number of CVs while choosing CVs that efficiently sample the desirable conformational space^44,45^: We thus chose two CVs - the N_α_–C_sp2_ distance as well as the α_BD_ value. α_BD_ was chosen in preference to α_FL_ because it varies over a wider range among the monomers evaluated (**Extended Data Fig. 6**).

### Metadynamics recapitulates relative reactivity of diverse sets of non-L-α-amino acid monomers

We began the metadynamics simulations with two initial conformations of each monomer. One conformation was identical to that used to initiate unbiased MD simulations (**Fig. 4** and **Extended Data Fig. 5**). The second conformation was altered by rotating the psi angle by 180°, to generate an alternative initial position of the nucleophile amine N_α_ relative to C_sp2_ of fMet. Examination of plots showing α_BD_ as a function of the N_α_–C_sp2_ distance after metadynamics simulations reveal minima that clearly differentiate highly reactive and less reactive monomers (**Fig. 5**). The metadynamics free energy surface (FES) contour plot of an RRM containing P-site fMet and A-site Met monomers shows excellent agreement with N_α_–C_sp2_ distance and α_BD_ values determined by cryo-EM for Met/Met. The global minimum N_α_–C_sp2_ distance is centered at 3.7 Å and that of α_BD_ is centered at 76°, while the fluctuations within 1 kcal/mol reach values between 3.4 - 4.4 Å and 61-90° respectively. These values show good agreement with both unbiased MD results (4 Å and 94°) and the metrics derived from the high-resolution structure reported here (3.3 Å and 93°). More importantly, the global minima for monomers that react efficiency within the PTC, notably pyridine **2** and (*1R,2S,*)-2-aminocyclopentane carboxylic acid **5**, populate conformations within the PTC with N_α_–C_sp2_ distances centered at or below 4 Å and α_BD_ between 90° and 105°. By contrast, monomers that are relatively inactive, notably *ortho*- and *meta*-aramids **3** and **4** and (*1S,2R*)-2-aminocyclopentane carboxylic acid **8** (the enantiomer of **5**) populate conformations with N_α_–C_sp2_ distances between 6 and 7 Å, albeit with α_BD_ centered again at 90°. The two moderately active 2-aminocyclopentane carboxylic acid isomers **6** and **7** populate conformational space defined either by long N_α_–C_sp2_ distances (**6**) or high α_BD_ values (**7**). These data suggest that the most reactive monomers significantly populate a conformational space characterized by an average N_α_–C_sp2_ distance of < 4 Å and an average α_BD_ of between 75 and 110°; low reactivity results when either of these boundary conditions are not met.

**Fig. 5.**
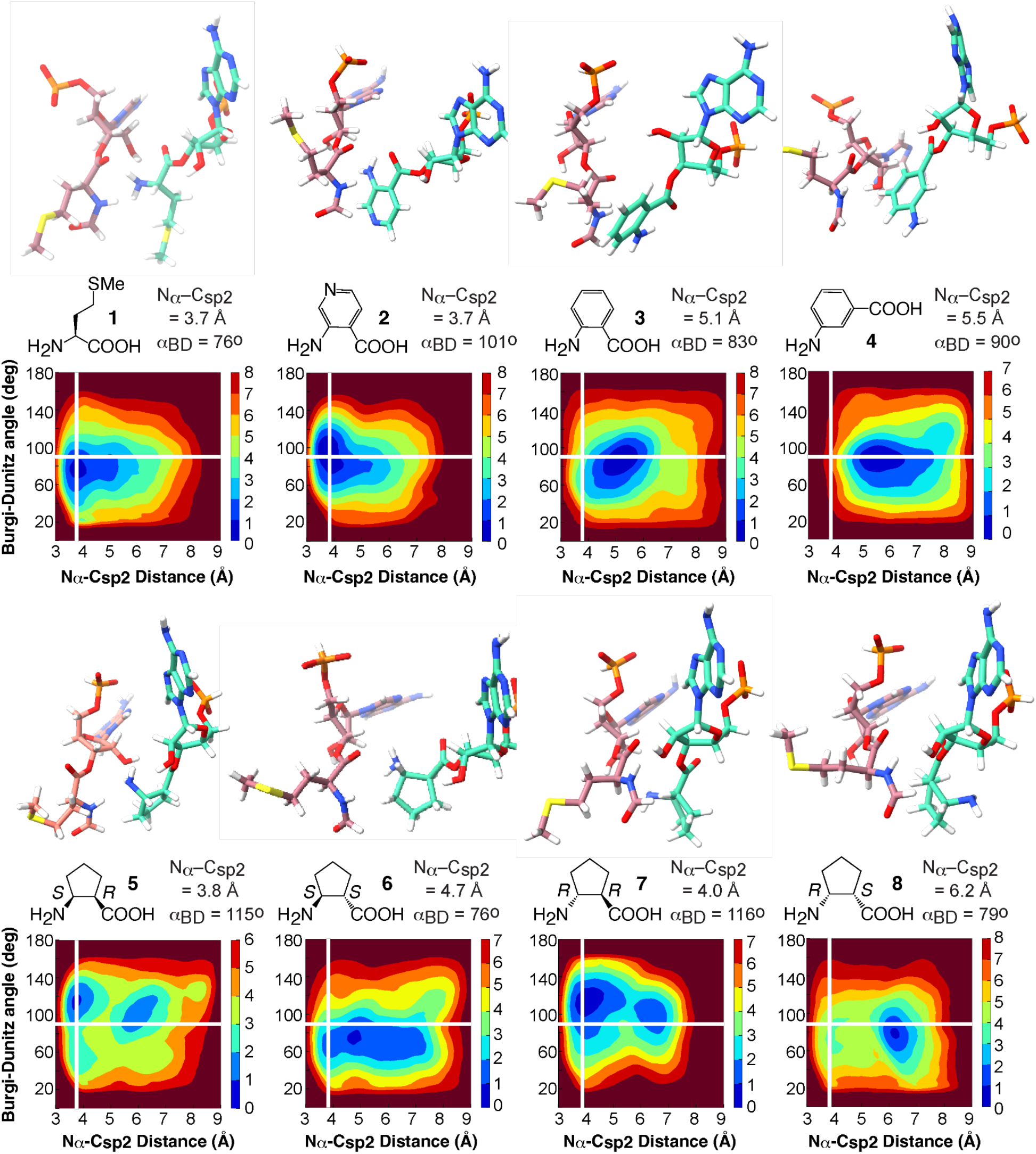
Plots of free energy surfaces from metadynamics simulations. Free energy surfaces (FES) (kcal/mol) of 30 Å RRM containing a P-site tRNA acylated with fMet and A-site tRNAs acylated with monomers **1-8** plotted along the Bürgi–Dunitz angle α_BD_ and N_α_–C_sp2_ distance collective variables. The metadynamics simulations that generated each FES began with the 2.1 Å cryo-EM model reported here but the A-site Met was *in silico* mutated to the other monomers while the P-site Met was converted to fMet. Each FES shown is the average of two metadynamics runs starting from orientations of the A-site monomers that differ by a 180° rotation about *psi*. The scale represents the free energy in kcal/mol where the global minima are set at 0 and therefore the various heights of the energy scales are based on the energetics of the fluctuations of the A-site monomers. The conformation and relative geometry of the P-site (rose) and A-site (green) monomers at the free energy minimum is shown above each plot.

Encouraged by these results, we next asked whether we could apply the metadynamics workflow to classify the relative reactivities of two structurally unrelated monomers that possess greater conformational freedom than those evaluated initially - notably (*S*)-β^3^-homophenylalanine **9** and (*S*)-β^3^-homophenylglycine **10**. These monomers acylate the identical tRNA with identical efficiencies in Flexizyme-promoted reactions, but (*S*)-β^3^-homophenylglycine **10** is significantly more reactive in *in vitro* translation reactions^46^. Application of the metadynamics workflow outlined above for monomers **2**-**8** to monomers **9** and **10** generated a pair of FESs that clearly differentiated the two monomers consistent with their reactivity (**Fig. 6**). The global minimum of the FES generated for an RRM containing monomer **10** in the A site mirrors that of reactive monomers **2** and **5**, notably a N_α_–C_sp2_ distance centered at or below 4 Å (3.7 Å) and α_BD_ of 111°. By contrast, the global minimum for monomer **9** mirrors that for inactive monomer **4,** notably a N_α_–C_sp2_ distance greater than 4.6 Å and a low α_BD_ of 72°.

**Fig. 6.**
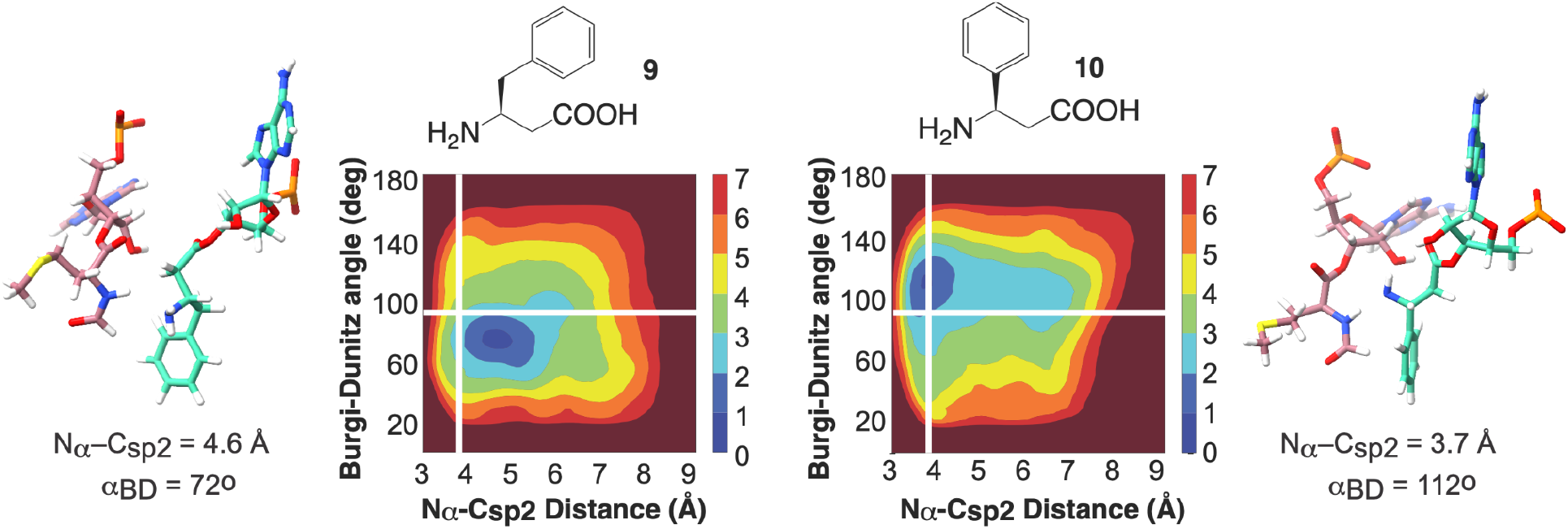
Metadynamics analysis of structurally unrelated monomers. The workflow accurately predicts the relative reactivities of two monomers that are structurally unrelated to monomers **2-8**, notably the less reactive (*S*)-β^3^-homophenylalanine **9** and the more reactive (*S*)- β^3^-homophenylglycine **10.** Shown are the free energy surfaces produced by the metadynamics workflow as well as the poses and geometries of the two systems at the global minimum.

In summary, here we develop a structure-informed, physics-based workflow to reliably classify the reactivity of non-L-α-amino acid monomers in amide bond forming reactions catalyzed by the *E. coli* ribosome. Non-L-α-amino acids monomers that populate a conformational space characterized by N_α_–C_sp2_ distances ≤ 4 Å and Bürgi-Dunitz angles ≥ 90° react efficiently within the PTC of wild type *E. coli* ribosomes. These predictions hold for three diverse families of molecules - aramids, β^3^-amino acids, and β^2,3^-cyclic amino acids. Monomers that cannot appreciably occupy a region in which the N_α_–C_sp2_ distance is less than 4 Å, even with an acceptable Bürgi-Dunitz angle, do not react. More broadly, the metadynamics workflow reported herein addresses two barriers impeding the ribosome-promoted biosynthesis of diverse heteroligomers. For applications *in vivo*, metadynamics can prioritize monomers for which orthogonal aminoacyl tRNA synthetase variants are needed. For applications *in vitro*, metadynamics can identify monomers that are likely to react within the PTC of wild type ribosomes, and those for which engineered ribosomes are needed.

## Methods

### Preparation of acylated tRNA

#### Synthesis of N-1 tRNA^fMet^

Templates for *in vitro* transcription were prepared using a double PCR amplification method. Base C1 was mutated to G for optimal T7 RNA polymerase initiation. First, long overlapping primers tRNA-fMet-C1G_temp F and RNA-fMet-C1G-A_temp R (**Extended Data Table 1**) were PCR amplified using Q5 DNA polymerase (NEB). Products were gel purified and amplified using short primers tRNA-fMet-C1G_amp F and tRNA-fMet-C1G-A_amp R (**Extended Data Table 1**). The 2nd base of each reverse primer in this step was modified with 2’OMe to prevent nonspecific addition by T7 RNA polymerase^47^. Amplification PCRs were 1x phenol chloroform extracted, 2x chloroform washed, and precipitated with 3x volumes of EtOH. DNA quantity was measured on an agarose gel using a known standard.

T7 *in vitro* transcription was performed in a buffer containing 50 mM Tris-HCl, pH 7.5, 15 mM MgCl_2_,5 mM dithiothreitol (DTT), 2 mM spermidine. Reactions contained 2.5 mM each NTP, 1:40 NEB murine RNase inhibitor, 25 μg T7 RNA polymerase (gift from Bruno Martinez, Cate lab), 0.0005 U/μL PPase, and ~ 1000 ng of DNA template per 100 μL of transcription reaction. Reactions were incubated for 16 h at 37 °C, treated with 1/20th volume RQ1 DNase (1 U/μL stock) for 30 min at 37 °C, and precipitated with 1/10th volume 3 M NaOAc pH 5.2 and 3x volumes EtOH. Precipitated tRNA was pelleted, washed once with 70% EtOH and resuspended in loading dye (95% formamide, 40 mM EDTA, 0.05% bromophenol blue, 0.05% xylene cyanol).

Gel purification was performed using ~ 20 cm long 12% polyacrylamide 1X TBE 7 M urea gels poured roughly 2 mm thick. Bands were excised using UV shadowing, crushed, frozen on dry ice briefly (with elution buffer), and eluted at 4 °C overnight in 300 mM NaOAc pH 5.2, 1 mM EDTA, 0.5% w/v SDS. Approximately 2 mL buffer was used per 500 μL transcription reaction. Eluted tRNA was pipetted off the gel debris and precipitated with 1 μL of glycoblue coprecipitant and 3x volumes of EtOH. Pelleted tRNA was resuspended in water and stored at −80 °C.

**Extended Data Table 1.**
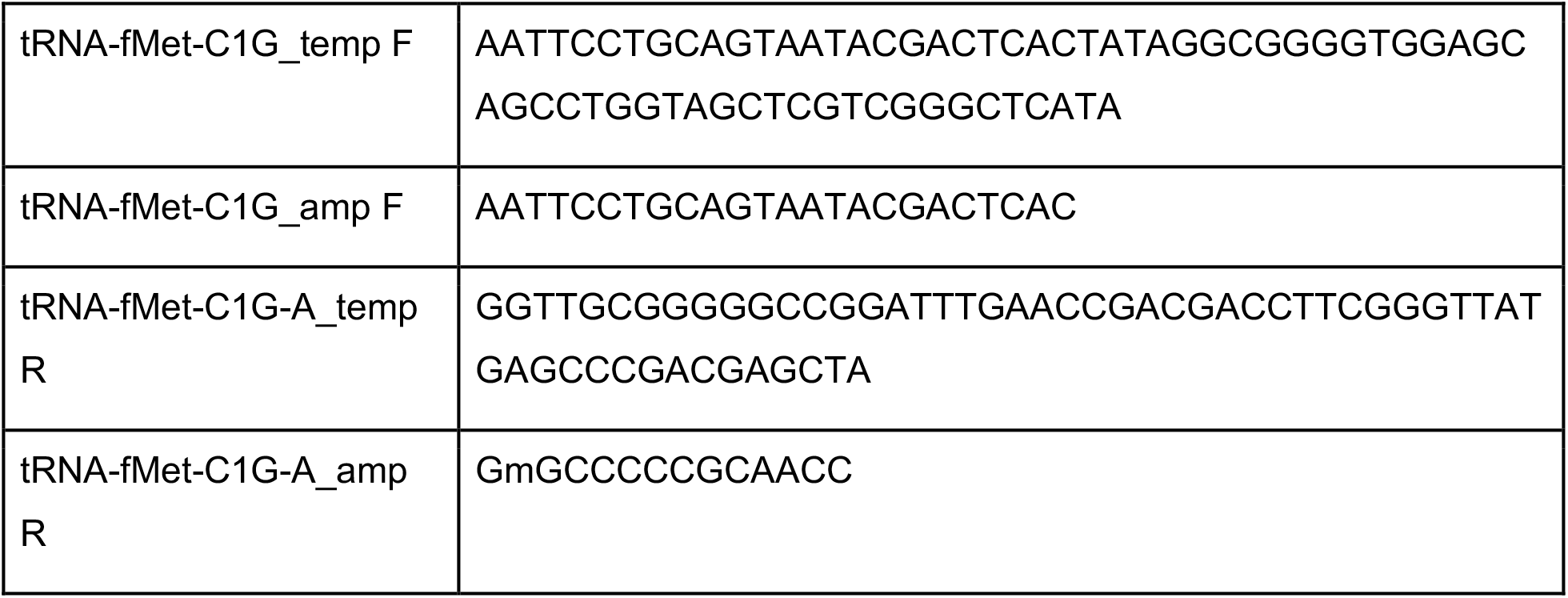
Primers used for tRNA template synthesis (‘mG’ indicates 2’ O-methylguanosine)

#### Purification of A. fulgidus CCA-adding enzyme

The full CCA-adding enzyme gene was purchased from Twist Bioscience and cloned into a plasmid carrying a T7 promoter and C-terminal 6x histidine tag. The resulting plasmid was transformed into BL21 (DE3) Rosetta2 pLysS cells for expression. 2x 1 L of LB broth + 100 μg/mL ampicillin was induced with a 1:100 dilution of overnight culture (grown at 37 °C). Cells were grown at 37 °C until OD_600_ was ~ 0.5 and induced with 0.5 mM IPTG for 3 h. Cells were pelleted and washed with lysis buffer (20 mM HEPES pH 7.5, 150 mM NaCl, 10 mM MgCl_2_, 1 mM DTT, 20 mM imidazole) and stored at −80 °C. For lysis, cells were resuspended in ~30 mL lysis buffer with a tablet of Pierce EDTA free protease inhibitor. Cells were lysed using a sonicator to deliver ~ 8000 J of energy and lysate was clarified at 18000 rpm in a JA-20 rotor (Beckman) for 30 min at 4 °C. Supernatant was applied to a 1 mL HisTrap column recirculating for ~ 30 min. The bound lysate was fractionated on an FPLC with 5 column volumes (CV) of lysis buffer and 10 CV of linear elution gradient from lysis buffer to lysis buffer with 500 mM imidazole. 0.5 mL fractions were collected and analyzed using SDS-PAGE. CCA-adding enzyme containing fractions were combined and concentrated in a 30000 Da MWCO spin filter and buffer exchanged into MS2-1 buffer. Protein was applied to a 1 mL heparin column and processed on an FPLC as previously described^48^. Protein-containing fractions were combined and dialyzed against 20 mM HEPES, 150 mM NaCl, 10 mM MgCl_2_, 1 mM DTT, 20% glycerol and stored at −80 °C. Final concentration was 54 μM in ~ 1 mL (extinction coefficient estimated 55350 M^-1^cm^-1^).

#### 3’NH_2_-ATP-tailing of N-1 tRNAs

The procedure used for tRNA amino-tailing was adapted from reference^49^. Equimolar amounts of N-1 tRNA and *A. fulgidus* CCA-adding enzyme (usually 2 μM) were combined in a reaction containing 100 mM glycine pH 9, 10 mM MgCl_2_, 1 mM DTT, 0.002 U/μL PPase, and 0.5 mM 2’-NH_2_-ATP (2’-amino-2’-deoxyadenosine-5’-triphosphate, purchased from Axxora). Reactions were incubated at 37 °C for 2 h. Degradation of tRNAs was seen if reactions were incubated past completion. NH_2_-tRNAs were extracted with 1:10 volume 3M NaOAc pH 5.2 and 1 volume acidic phenol chloroform, cleaned twice with 1 volume chloroform, and precipitated with 3 volumes EtOH. NH_2_-tRNAs were resuspended in water and reaction yield was analyzed on a 10% acrylamide 7 M urea TBE gel (20 cm). 1-2 pmol of tRNA was loaded per lane and the gel was stained with Sybr Green II after running. NH_2_-tRNAs were stored in aliquots at −80 °C.

#### Purification of MetRS

6xHis-tagged MetRS expression plasmid of was a gift from Patrick Ginther (Schepartz Lab, UC Berkeley). The plasmid was transformed into BL21 (DE3) Codon+ RIL cells (T7 promoter). Culture overnights were diluted into ZYM-5052 autoinducing media^50^ and expressed overnight at 37 °C. Cells were pelleted and resuspended in lysis buffer (20 mM Tris pH 7.8, 150 mM NaCl, 5 mM imidazole, 0.5 mM EDTA) (~ 35 mL). Cells were lysed with a sonicator on ice until ~ 8000 J had been delivered to the sample. Lysate was clarified by centrifugation at 18000 rpm (JA-20 rotor; Beckman) at 4 °C for 30 min. Supernatant was applied to a 5 mL HisTrap column and the column was attached to an FPLC. Protein was purified on the FPLC by washing with 5 CV of lysis buffer with 23 mM imidazole and eluting with a linear gradient of 20 CV from 23-500 mM imidazole. Fractions containing the desired protein were pooled, dialyzed overnight against 50 mM HEPES pH 7.5, 100 mM KCl, 10 mM MgCl_2_,7 mM BME, 30% glycerol, and concentrated in spin filters. Protein was stored at −80 °C in aliquots.

#### Aminoacylation of tRNAs

All aminoacylation reactions were performed in a buffer containing 50 mM HEPES pH 7.5, 20 mM MgCl_2_, 10 mM KCl, 2 mM DTT, 10 mM ATP, 1:40 volume RNase inhibitor (murine, NEB), 5-10 mM amino acid. Aminoacylation enzymes were usually used at 1 μM and tRNAs at 2-5 μM depending on stock concentration. Generally, a 1:5 enzyme:tRNA ratio was not exceeded. Reactions were incubated at 37 °C for 30 min and then extracted and precipitated as in *“3’NH2-ATP-tailing of N-1 tRNAs.”*

### 70S ribosome preparation

#### Met-Met ribosome complex formation

All steps toward the Met-Met ribosome structure were performed essentially as described^28^. Briefly, the complex was formed by incubating 100 nM 70S ribosomes, 5 μM mRNA, ~ 1.6 μM Met-NH-tRNA^fMet^, and 100 μM paromomycin in buffer AC (20 mM Tris pH 7.5, 100 mM NH_4_Cl, 15 MgCl_2_, 0.5 mM EDTA, 2 mM DTT, 2 mM spermidine, 0.05 mM spermine) for 30 min at 37 °C. The mRNA sequence GUAUAA**GGAGG**UAAAA<u>AUGAUG</u>UAACUA was synthesized by IDT. (Shine-Dalgarno sequence in bold, 2x AUG codons underlined.)

### Cryo-EM sample preparation

The sample was prepared for imaging on 300 mesh R1.2/1.3 UltrAuFoil grids with an additional layer of float-transferred amorphous carbon support film. The grids were washed in chloroform prior to carbon floating. Before applying the sample, grids were glow discharged in a PELCO easiGlow at 0.39 mBar and 25 mAmp for 12 seconds. 4 μL of sample was deposited onto each grid and left for 1 minute. The grid was then washed with a buffer containing 20 mM Tris pH 7.5, 20 mM NH_4_Cl, 15 MgCl_2_, 0.5 mM EDTA, 2 mM DTT, 2 mM spermidine, and 0.05 mM spermidine by successively touching to three 100 μL drops of the buffer immediately prior to freezing. Grids were blotted and plunge-frozen in liquid ethane with an FEI Mark IV Vitrobot using the following settings: 4°C, 100% humidity, blot force 6, blot time 3. Grids were clipped for autoloading and stored in LN_2_.

### Cryo-EM Data Collection

Cryo-EM data collection parameters are summarized in **Extended Data Table 2**. Dose-fractionated movies were collected on a Titan Krios G3i microscope at an accelerating voltage of 300 kV and with a BIO Quantum energy filter. Movies were recorded on a GATAN K3 direct electron detector operated in CDS mode. A total dose of 40 e^-^/Å^2^ was split over 40 frames per movie. The magnification was 102,519 for a physical pixel size of 0.8296 Å and super-resolution pixel size of 0.4148 Å (based on pixel size calibration performed after the final structure was obtained). Data collection was automated with SerialEM^51^, which was also used for astigmatism correction by CTF and coma-free alignment by CTF. One movie per hole was collected using stage shift to move between center holes of a 3×3 hole template and image shift to collect on the surrounding 8 holes. The defocus ramp was set to range between −0.5 and −2 μm.

### Image Processing

All RELION steps were performed in RELION 3.1^52^. Motion correction was performed with MotionCor2^53^ within the RELION GUI and micrographs were binned to the physical pixel size (believed to be 0.81 Å at the outset). CTF parameters were estimated using CTFFind4^54^. Micrographs with poorly fitting CTF estimates were rejected based on visual inspection, leaving 6673 movies for processing. Particle auto-picking was performed using the Laplacian-of-Gaussian method in RELION, yielding 1,021,926 particles. These were then extracted from the micrographs with rescaling to ⅛ the full size and underwent 3 rounds of 2D classification. The 735,114 particles from the good 2D classes were then re-extracted at ¼ the full size and imported into cryoSPARC^55^ v.3.1.0 for heterogeneous refinement with 6 volume classes. The reference ribosome volume for this job was generated from 1VY4 PDB coordinates^22^ in EMAN2^56^. Two of the resulting classes corresponded to clean 70S volumes, so these classes were exported back to RELION and pooled (513,480 particles).

To separate out classes with different positions/rotation of the small subunit relative to the large subunit, 50S-focused refinement was first performed on all the particles, followed by 3D classification without alignment into 5 volume classes. In addition to subtle shifts in the 30S subunit, this step is useful for removal of particles in minor states like those containing E-site tRNAs only, for example. From there, further pooling and focused classification on A- and E-site tRNAs (without alignment) were performed, details of which are summarized in **Extended Data Fig. 3a**. As all particles that were accepted at this stage contained P-site tRNAs, and the A and E sites appeared to produce more meaningful separation, classification on the P site was not pursued.

For the final structure, all particles in the three classes selected at the 30S rotation stage were extracted at full size and pooled for 50S-focused refinements prior to and in between CTF refinement^57^, Bayesian Polishing^58^, and CTF refinement again. This was to take advantage of high resolution from the well-ordered large subunit and maximize particle number for optimum performance of these jobs, and the resulting post-processed volume was also used for pixel size calibration. Particle classes of interest that had been identified earlier were separated again for final 50S-focused refinements using python scripting to pull the appropriate subsets from the polished particle metadata.

From the classification procedure, there were 3 initial A-site classes that were examined individually as well as merged, and with and without further sorting on the E site. Ultimately, variation across maps indicated some residual disorder in fine details of the substrates, with some features alternately improved or worsened in different maps, and there was no single map representing the best set of features across A- and P-site Met residues or their surroundings (**Extended Data Fig. 3b**). Taken together, it is clear that we are still somewhat limited by inherent dynamics of substrates in the active site. Additionally, the line between meaningfully distinctive classes or conformations and continuous motions of the complex(es) presents a challenge, particularly when particle number starts to become limiting. A merged map of two of the A site classes was ultimately chosen for modeling, containing 129,455 particles and with a global resolution of the large subunit and tRNAs going to 2.1 Å resolution.

### Pixel Size Calibration

The pixel size was calibrated in Chimera^59^ using the “Fit to Map” function and the high resolution pooled-particle map against the 50S subunit coordinates from the X-ray crystal structure PDB 4YBB^60^. The best cross-correlation value was obtained at pixel size 0.8296 Å. Half-maps for our refined volume were rescaled to the calibrated pixel size in Chimera and post-processing was performed with these half-maps in RELION to obtain the FSC curve and final resolution estimate (**Extended Data Fig. 2**).

### Modeling

PDB 7K00^28^ was taken as a starting model. This model contained tRNA^fMet^ in the P site and tRNA^Val^ in the A site, whereas only tRNA^fMet^ was used in the current work, so the A-site tRNA was replaced. The 30S subunit coordinates were not used as our focus was on the PTC. Real-space refinement of the coordinates was performed in PHENIX^61^ and further adjustments to the model were done manually in Coot^62^, mainly around the PTC. Further additions to the model included Mg^2+^ and K^+^ ions, water molecules, and spermidine ligands near the tRNA CCA ends, as well as residues 2-7 of r-protein bL27. The linkage between A- and P-site tRNAs and Met monomers is modeled as an amide linkage to reflect experimental conditions for structure determination. Map-vs.-model FSC was calculated in PHENIX^61^ (**Extended Data Fig. 2b**). Model refinement statistics are summarized in **Extended Data Table 2**.

### Molecular dynamics simulations

As described in Results, the starting point for our MD simulations was the RRM, which involved rigid-body docking of ordered Mg^2+^ ions from PDB 7K00 into preliminary coordinates for the cryo-EM structure reported here, which ultimately included additional ions not present in 7K00. It should be noted that static structures fail to capture the dynamic interactions between the RNA and ions^63^. Moreover, monovalent cations generally interact with RNA as highly dynamic species, creating a diffusive ion environment, while divalent cations, such as Mg^2+^ form an ion-hydration shell^64,65,26^. Hence, the rigid-body docking of the Mg^2+^ ions from 7K00 described above was considered sufficient for our unbiased and biased MD calculations.

The resulting structure was solvated using the simple point charge (SPC) water model^66^. K+ and Cl- ions corresponding to 0.15 M concentration were added as well as K+ counterions to neutralize the system. The final simulation box measured 95 Å along each side and consisted of ~88000 atoms. The OPLS4 force field^67^ and Desmond MD system (Schrödinger Release 2022-2) as implemented within Schrödinger Suite (release 2022-2) were used in this study. For fMet and all the non-L-α-amino acids monomers, Force Field Builder (Schrödinger release 2022-2)^67^ was used to optimize the missing torsions, which does so by fitting the molecular mechanics torsional profiles to those obtained based on quantum mechanics calculations.

The systems were initially minimized and equilibrated with restraints on all solute heavy atoms, followed by production runs with all but the outer 10 Å C1’ and Cα atoms unrestrained. The NPT ensemble was used with constant temperature at 300K and Langevin dynamics. The production runs were carried out for 300ns in triplicates, changing the initial velocity seeds for each run. The conformational analysis, including the distances and angles, was calculated using Schrödinger’s Python API (Schrödinger Release 2022-2). The representative pose for the test fMet-Met run (**Fig. 2c**) was generated using the trajectory RMSD-based clustering method^68^ as implemented in Maestro. The RDF was generated using Schrödinger’s built-in RDF panel.

### Metadynamics

Desmond^69^ (Schrödinger Release 2022-2) was used for the metadynamics runs. The equilibration stage was the same as for the MD runs above and the metadynamics production runs were carried out in duplicates (starting from different conformations as outlined above) for 100 ns each. The N_α_–C_sp2_ distance and the Bürgi–Dunitz angle were used collective variables (CVs). The biasing Gaussian potential (“hill”) of 0.01 kcal/mol was used and the width of 0.15 Å for the N_α_–C_sp2_ distance and 2.5° for the Bürgi–Dunitz angle α_BD_ were applied. Analysis of the runs were performed with Schrödinger’s Python API as well as *in-house* Python scripts.

## Data availability

The cryo-EM map and model reported here are deposited at the EMDataBank and RCSB Protein Databank with accession codes EMD-XXXXX and PDB XXXX. Additional data supporting the findings of this manuscript are available from the corresponding authors upon reasonable request.

## Code Availability

Python scripts written using Schrödinger’s API for the analysis of the data from the MD and Metadynamics simulations are available upon request.

## Acknowledgements

This work was supported by the NSF Center for Genetically Encoded Materials (C-GEM), CHE 2002182. We are grateful to Dr. Sarah Smaga and Dr. Josh Walker for insights regarding monomer selection, and to Dr. Matt Repasky (Schrödinger) and members of the Cate and Schepartz labs for comments on the manuscript. We also thank Dr. Daniel Toso and Paul Tobias for assistance with cryo-EM data acquisition and management.

## Contributions

S.J.M., J.H.D.C., A.S., and A.M.A. conceived the study. A.M.A., Z.L.W., I.K., J.H.D.C., S.J.M. and A.S. designed the project. A.M.A. led computational experiments and analysis. Z.L.W. led cryo-EM experiments and analysis. F.R.W. prepared ribosomal complexes for cryo-EM. A.M.A., Z.L.W., I.K., J.H.D.C., S.J.M. and A.S. analyzed and interpreted data. Z.L.W., I.K., S.J.M., J.H.D.C., A.S. and A.M.A. prepared the manuscript.

## Supplementary Information

### Extended Data Figs. 1–6, Tables 1-2, references

**Extended Data Fig. 1.**
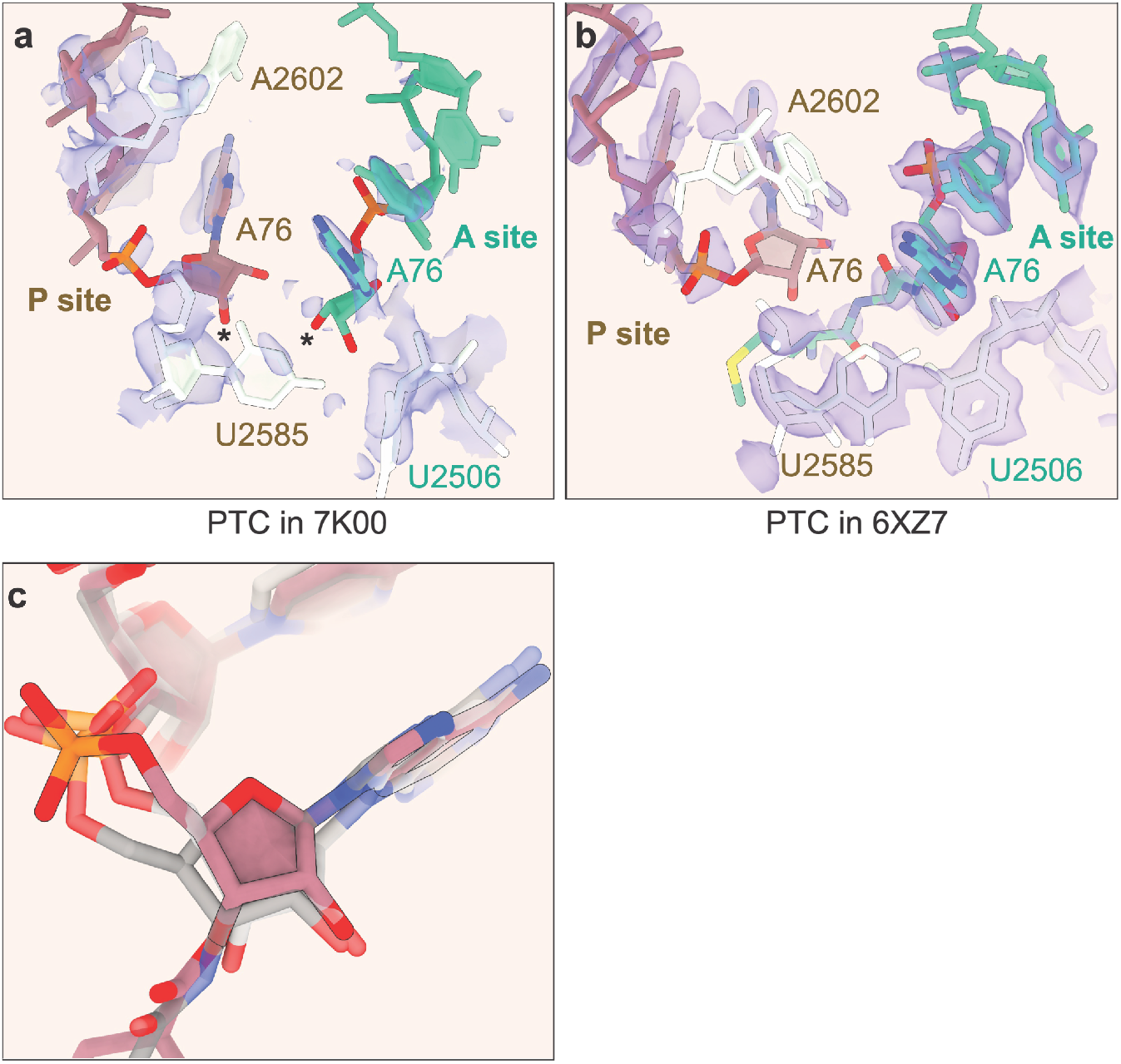
Details in the PTC of prior structures. **a,** Close-up of the PTC in PDB 7K00 illustrates the poor quality of local density in this map (indigo) around the P site (tan) and A site (green) tRNAs. Asterisks indicate A76 O3’ atoms which would bear a monomer. PTC bases U2506, U2585, and A2602 of the 23S rRNA are shown in white. The deposited 50S- focused map is shown with an additional B-factor of 30 Å^2^ applied, supersampled for smoothness. **b,** Close-up of the PTC in PDB 6XZ7, showing dipeptide in the A site (green with atomistic coloring) and uncharged P-site tRNA (tan). The map has been supersampled for smoothness. Contour levels for maps shown in both **a** and **b** are chosen to illustrate disorder relative to better-resolved map regions, but lower contour levels and/or higher B-factor blurring brings out features supporting models. **c,** P-site tRNA A76 backbone conformation in our model (pink, black outline) compared to 6XZ7 (light gray) and 1VY4 (dark gray).

**Extended Data Fig. 2.**
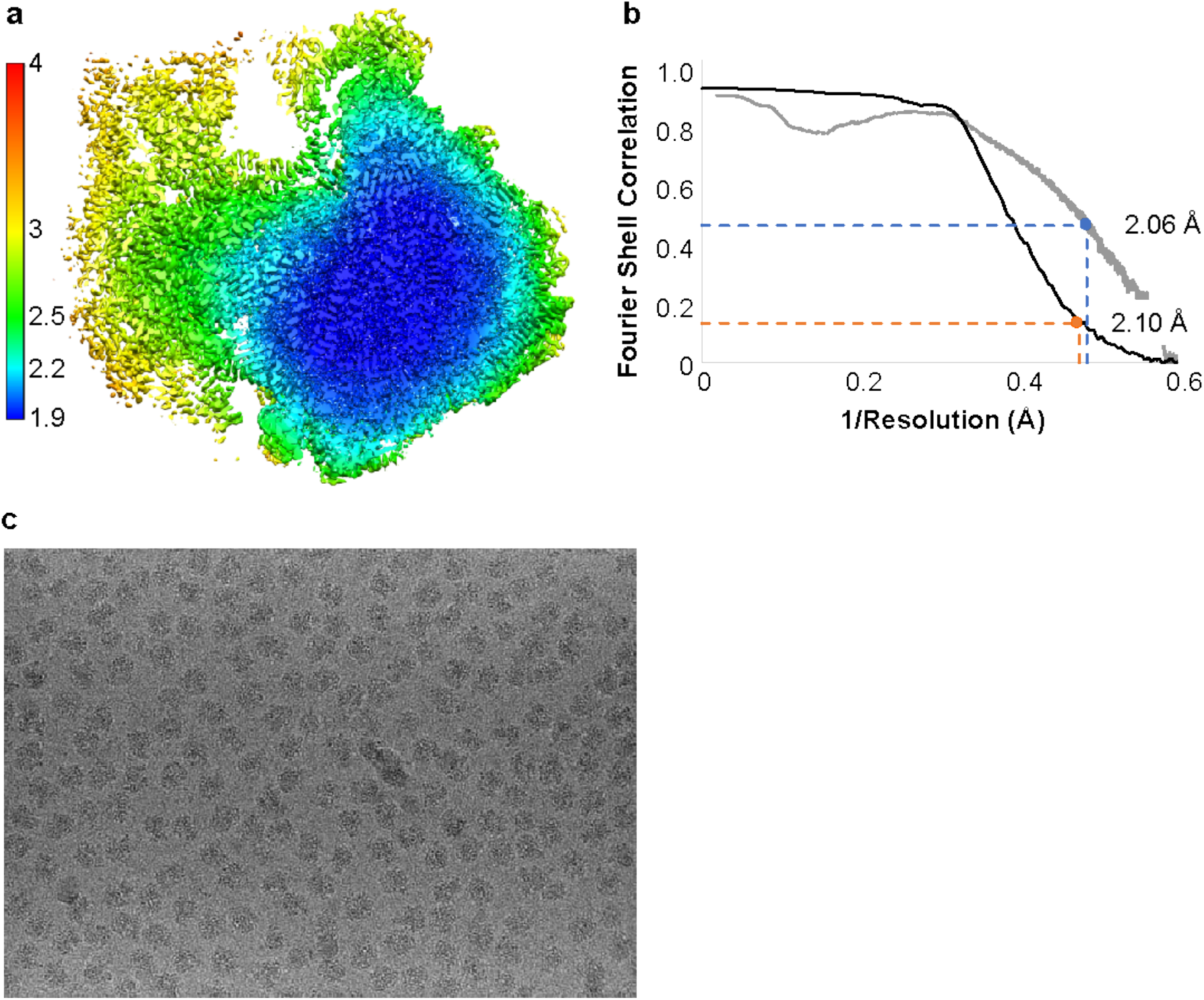
Resolution of the 50S subunit in the Met-Met ribosome structure. **a,** Local resolution of the 50S-focused cryo-EM map, **b,** FSC curves indicating resolution of the 50S subunit. Half-map FSC (black) is at 2.10 Å at the gold-standard cutoff value 0.143; map-vs.- model FSC (gray) is at 2.06 Å at the 0.5 cutoff. Only the large subunit was masked for the halfmap FSC, and **c,** representative micrograph showing individual ribosome particles. The micrograph was low-pass filtered to 20 Å for easier visualization of particles.

**Extended Data Figure 3.**
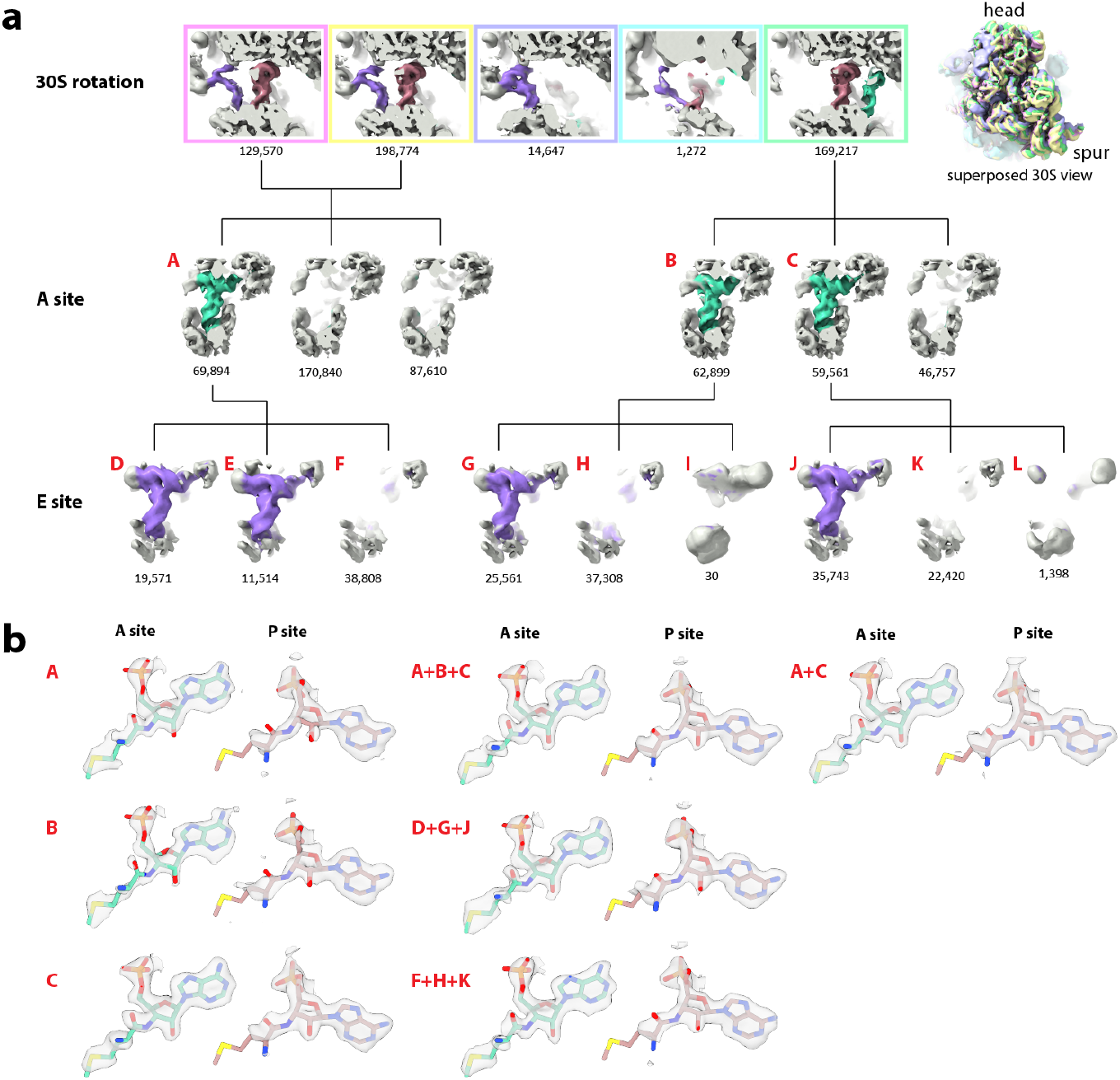
**a,** Flowchart illustrating classification workflow applied during cryo-EM data processing. Apart from the top right 30S view, density for A-site, P-site, and E-site tRNAs are shown in green, pink, and purple, respectively, with surrounding ribosomal density in gray. The number of particles in each class is shown beneath its image and all classes with A-site tRNA are identified with an uppercase letter in red. Top row: After an initial consensus 50S-focused refinement, particles were first sorted into 5 classes to discriminate between populations with different relative 30S subunit positions. Pink, yellow, purple, cyan, and green outlines on the zoomed-in cutaway panels are colored to match corresponding superposed whole ribosome maps on the top right, showing subtle rotations of the small subunit. Pink- and yellow-outlined classes were merged based on similarity. Middle row: particles were then classified on the A site. Bottom row: particle classes with A-site tRNA were further classified on the E site. Interestingly, only one of the A-site classes included particles with E-site tRNA in two different positions (classes D and E), however, this did not affect analysis of PTC density. **b,** Close-ups of A- and P-site density for select reconstructions that were compared to be chosen for modeling. While a number of permutations were considered, here we show the initial classes A, B, and C before sorting on the E site, as well as A+B+C and A+C merged. We also show D+G+J merged and F+H+K merged, representing the most homogeneous sets of particles in terms of tRNA occupancy. However, classes A+C were chosen for modeling the PTC. Cryo-EM density is shown in gray.

**Extended Data Fig. 4.**
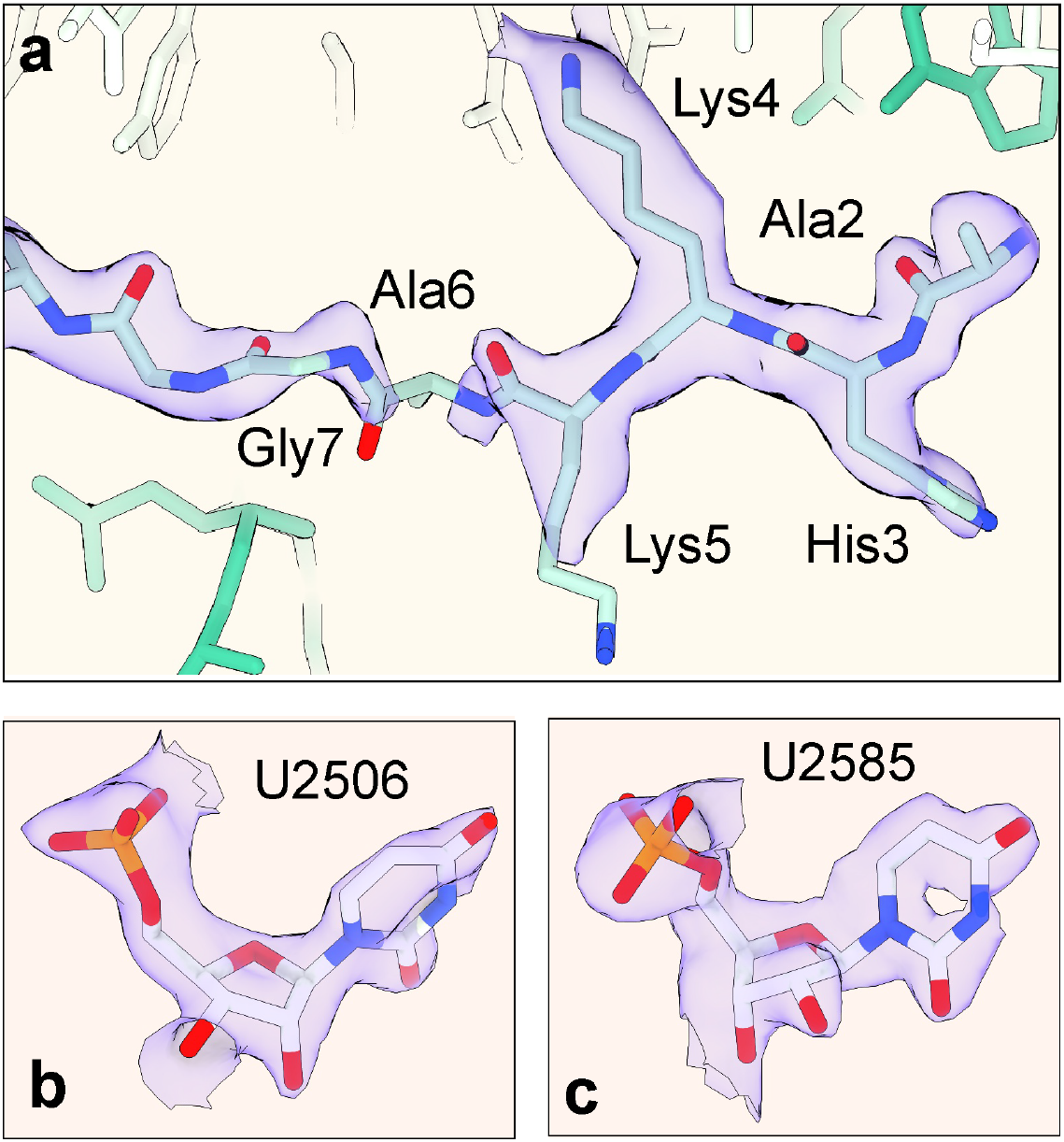
Structural model of the N-terminus of ribosomal protein bL27. **a**; Newly modeled section of r-protein bL27 (Ala-2 to Gly-7) (light blue). 23S rRNA is shown in white and A-site tRNA in green, with EM density shown in indigo. The map here has been postprocessed with a B-factor of 10 Å^2^. **b, c,** bases U2506 and U2585 modeled in the cryo-EM density. The map here has been post-processed with a B- factor of −13 Å^2^ and supersampled for smoothness.

**Extended Data Fig. 5.**
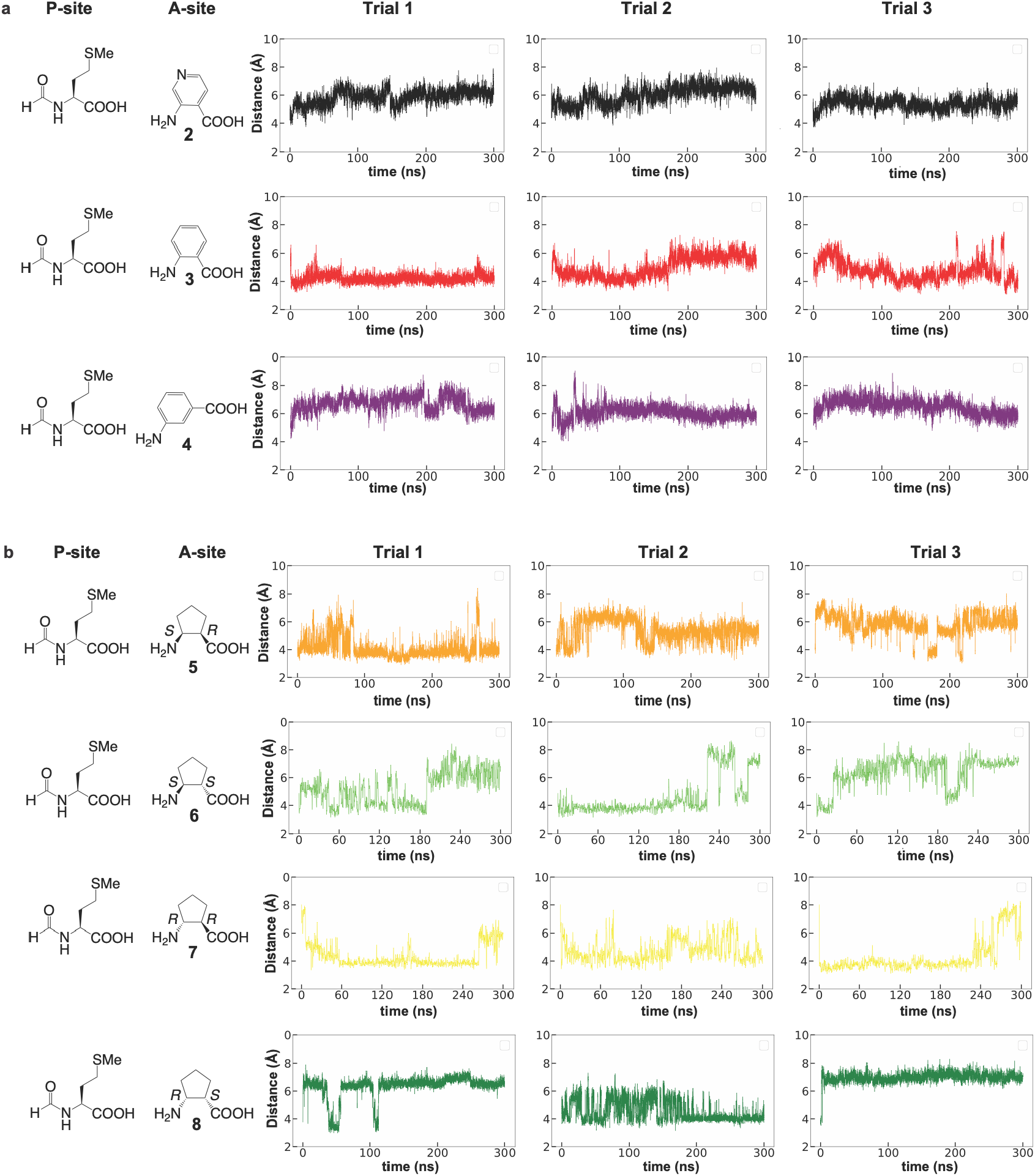
Trajectories from MD simulations of different monomers in the A site. Trajectories illustrating evolution of the distance between the A-site nucleophile (N_α_) and the P-site fMet carbonyl electrophile (C_sp2_) over 300 ns within the 30 Å Reduced Ribosome Model (RRM). **a,** aramids **2-4**; **b,** β^2,3^-2-aminocyclopentane-1-carboxylic acids **5-8**. We used Schrödinger’s Force Field Builder (Schrödinger release 2022-1) to parametrize the existing OPLS4 force field^67^ to accurately describe the conformational dynamics of each new monomer. Maestro’s 3D Builder was then used to assemble the new ester bond and minimize the system to release clashes with surrounding residues.

**Extended Data Fig. 6.**
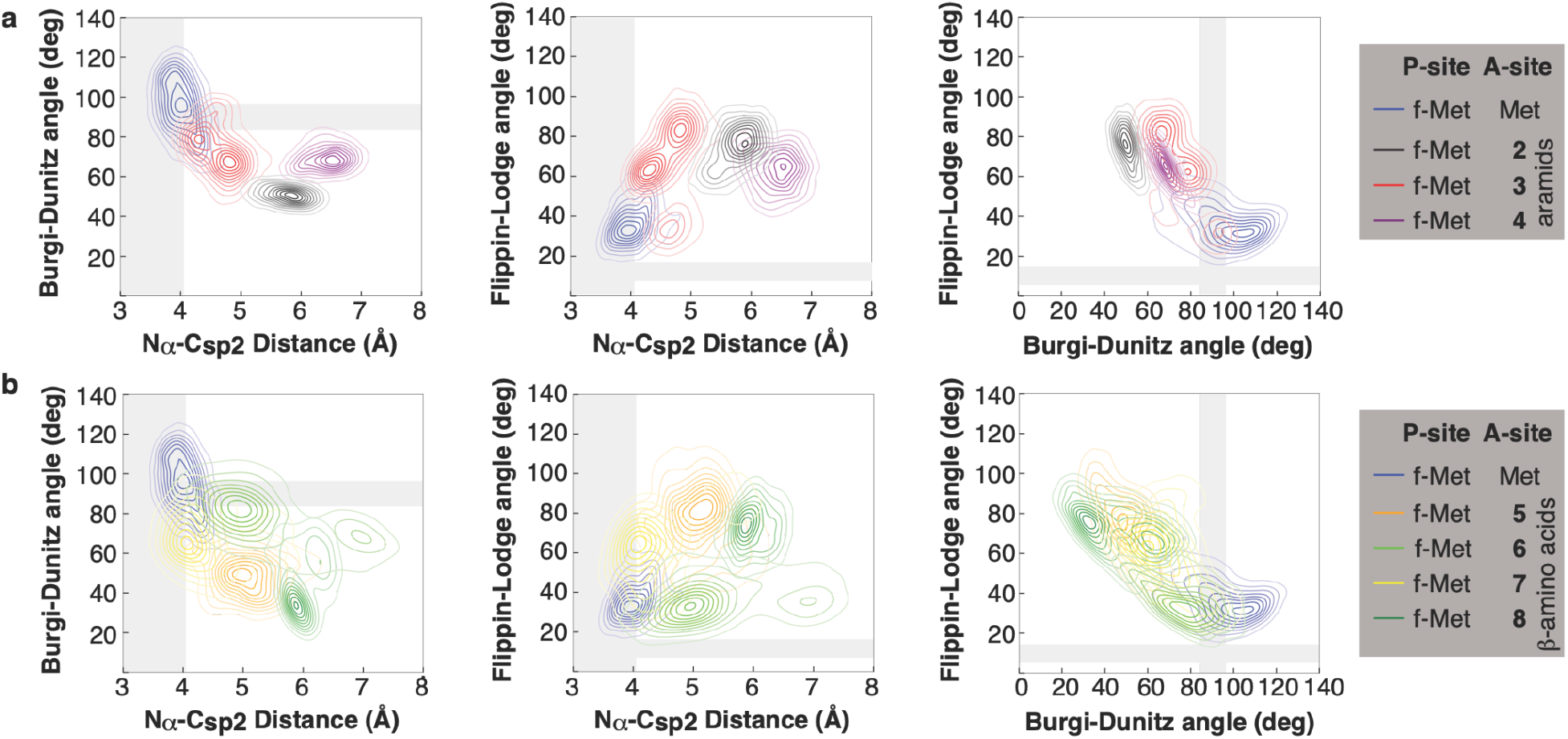
Scatter plots of the three geometric measurements used to evaluate conformational fluctuations. **a,** fluctuations of aramids **2-4** and **b, fluctuations of** cyclic β^2,3^-amino acid 2-aminocyclopentane-1-carboxylic acid monomers **5-8** in comparison to Met **1** in the PTC of RRM throughout the MD runs are shown. Specifically, the N_α_–C_sp2_ distance is plotted against either the Bürgi–Dunitz angle α_BD_ (left) or the Flippin-Lodge angle (center), and the two angles are plotted against each other (right). The gray zones highlight N_α_ - C_sp2_ distances of between 3 and 4 Å, and values of α_BD_ and α_FL_ of between 90° and 105° and 5° to 20°, respectively.

**Extended Data Table 2.**
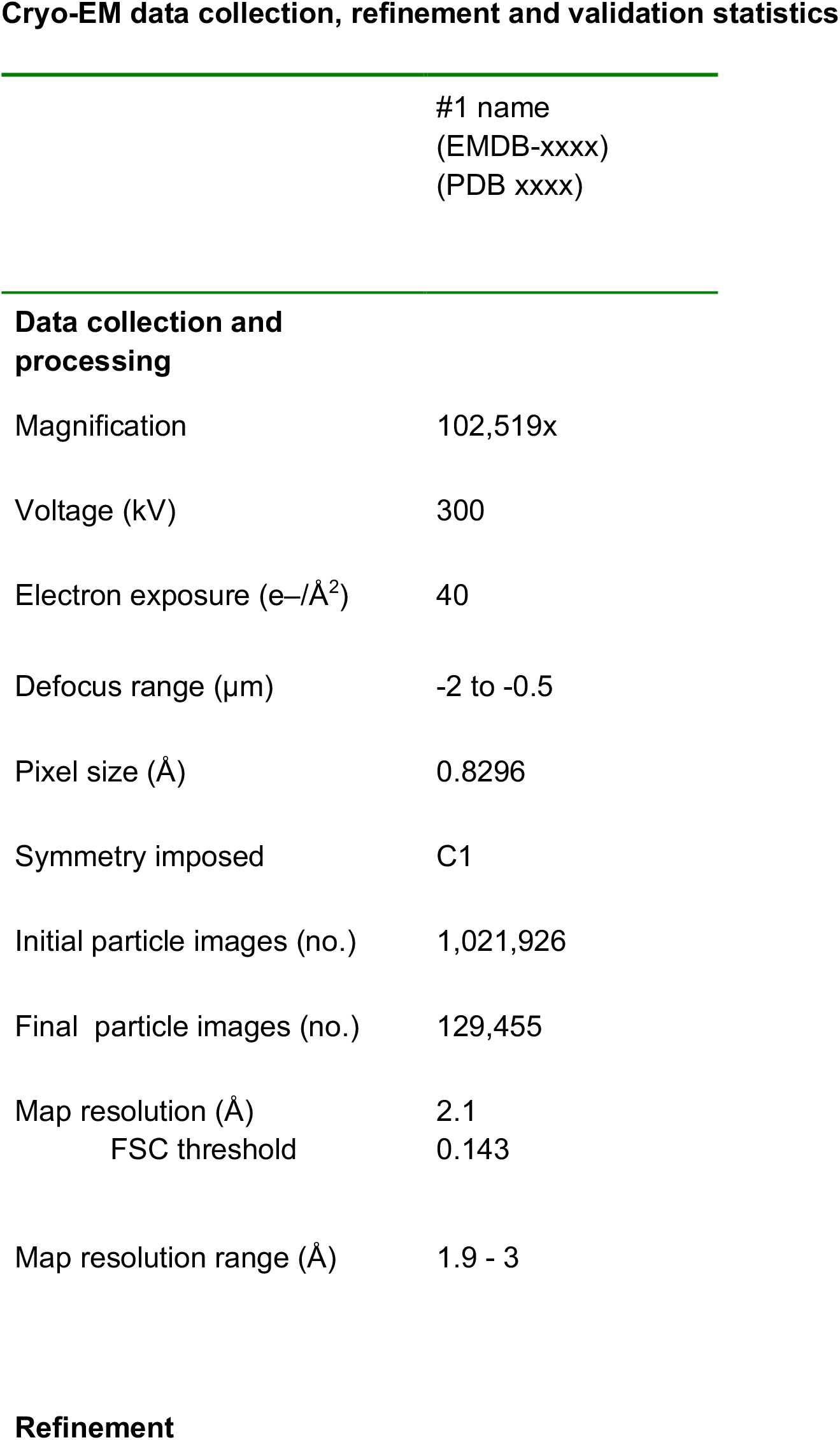
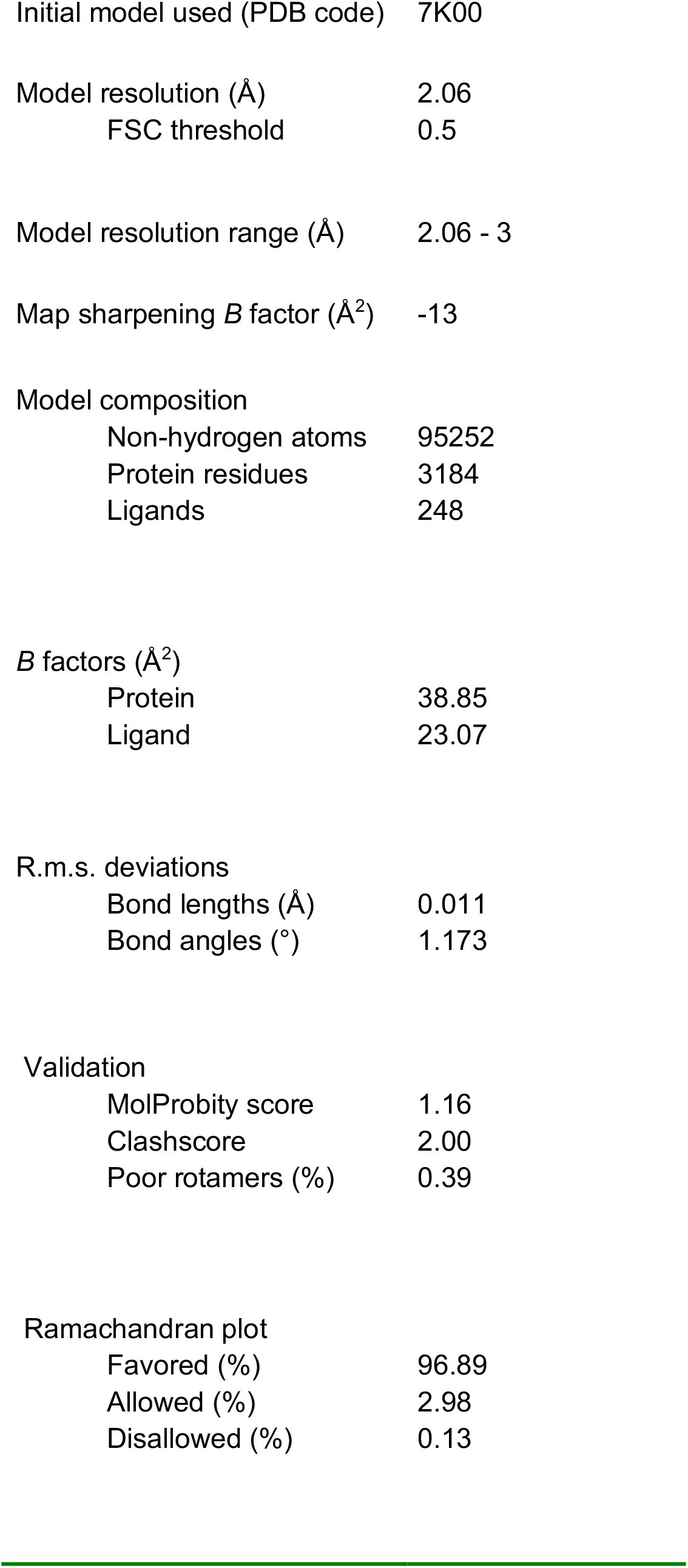
Cryo-EM data collection parameters and Refinement and validation statistics

